# A binocular perception deficit characterizes prey pursuit in developing mice

**DOI:** 10.1101/2022.05.18.492552

**Authors:** Kelsey Allen, Rocio Gonzalez-Olvera, Milen Kumar, Ting Feng, Simon Pieraut, Jennifer L. Hoy

## Abstract

The mouse continues to be one of the most powerful models in which to address the molecular and cellular mechanisms underlying mammalian visual system development. In particular, integration of binocular information at the cellular level has long been studied in specific regions of the mouse visual cortex to gain mechanistic insight into the development of vision. However, we lack an understanding of the development of the visual perceptions themselves in mice and other species that might rely on successful binocular integration. Further, the superior colliculus also processes binocular input and it remains unclear how early visual experience differentially impacts activity in this brain area during behavior. To address these major outstanding questions, we quantified the natural visually-guided behavior of postnatal day 21 (P21) and adult mice using a live prey capture assay and a computerized-spontaneous perception of visual objects tasks (C-SPOT). Analysis of both behavioral assays revealed robust and specific binocular visual field processing deficits in P21 mice as compared to adults. In addition, c-Fos expression in the anterior region of the superior colliculus (SC), the region that would process stimuli located in the binocular visual field, was highly different between P21 mice and adults after C-SPOT. We thus exploited a natural visual pursuit behavior and C-SPOT to provide the first demonstration of a specific visual perception deficit related to binocular integration in developing mice.

**Highlights:**

- Juvenile (P21) mice robustly investigate live insects
- Insect pursuit behavior relying on binocular vision is immature in P21 mice
- Visually-induced arrest responses are similar between P21 and adult mice.
- Ethologically-relevant visual experience differentially increases c-Fos expression in the superior colliculus of juveniles versus adults.

## Introduction

Mice are used to study many aspects of visual system development and have significantly advanced our understanding of how visual stimulus encoding in the brain matures (see^1–4^ for representative reviews). This is particularly true regarding our understanding of how and when neurons in the primary visual cortex acquire their ability to integrate binocular input^4^. However, how developmental changes in visual stimulus encoding such as binocular integration relates to age-specific natural visual perception and behavior remain unexplored^5,6^. Further, the superior colliculus (SC) is capable of representing a rich repertoire of binocular information and controlling many important visual behaviors in the adult mouse^7,8,9^. It is also unknown whether and how visual experience may shape the emergence of many receptive properties including binocular processing in the SC and how such changes could influence visual behavior, and vice versa, over development in the mouse^8,10^. Thus, generating an understanding of these two aspects of mouse visual development, age-characteristic visual behavior and the role of the superior colliculus in controlling those visual behaviors, are critical to linking a large body of existing molecular and neural circuit work to their ultimate implications in understanding plasticity in visual perception.

Studying the development of innate visual behaviors is a particularly fruitful approach to identifying the causal relationship between highly conserved visual behavior and perception and the structural and functional refinement of visual neural circuits^11,12^. In particular, the complex visual processing required for mice, other rodents and even primates to capture prey is an ideal natural context to investigate fundamental and conserved functions of the visual system and its development^13–15^. Key behaviors and perceptions deployed during this evolutionarily important behavior include object identification, search and pursuit^13,15,16^. Pursuit especially encompassing complex visual control of motor movements that require spatial precision and predictive coding to yield desired results^15,17,18^. There is also evidence that juveniles must have adequate time to practice the behaviors needed for prey capture in order to survive in the wild^19,20^. Larval zebrafish also demonstrate remarkable plasticity in prey capture behavior that requires increasing integration between the tectum (homologous to the SC) and forebrain circuitry^21^.

Similarly, adult mice significantly change their responsiveness to virtual “prey-like” stimuli presented in our computerized-spontaneous perception of visual objects tasks (C-SPOT) after having previous experience with live crickets^22^. Taken together, this emerging body of work supports the idea that studying the development of visually-guided prey capture in the mouse offers a unique and powerful paradigm to mechanistically decipher plasticity in visual behavior at the molecular and circuit level.

Of note, it has been recently established that binocular vision and sensory processing in the superior colliculus are required for optimal visually-guided prey capture in the mouse^23–25^. However, it remained unknown whether juvenile mice exhibited insect hunting, or hunting-like behavior, and whether binocular-dependent perceptions and/or prey capture-related activity in the superior colliculus might also be developmentally regulated in the mouse. In this study, we addressed these questions by assaying both live prey capture behavior and assaying c-Fos expression in the SC after visually-guided responses to prey-like virtual objects in our C-SPOT assay at P21 versus mature adults. Our analyses revealed that P21 mice lack a prominent binocular visual field bias that enhances successful “target” approach and pursuit in the adult in both the live and virtual behavioral assays. Additionally, C-SPOT revealed that visually-guided approach behavior was specifically different in early development relative to visually-induced arrest behavior driven most frequently by small moving objects present in the monocular visual field. Analysis of c-Fos^26^ expression in the SC after C-SPOT revealed a distinct pattern of enhanced cellular activity in the anterior regions of the SC in juveniles responding to “standardized” visual motion stimuli relative to the adults. Overall, our findings provide unique insight into the relationship between binocular vision used in a natural context by a mouse and its development and implicate experience-dependent changes in the anterior SC as relevant to understanding the maturation of binocular perception. Finally, we demonstrate that C-SPOT can readily probe the mechanisms underlying the development of binocular and other aspects of vision as well as assay mouse models of neurodevelopmental disease that impact vision and visual system plasticity^27^.

## Results

To understand how the mouse relates to other species where the young must practice prey capture behavior^19^, we first determined whether postnatal day 21 (P21) mice, before weaning and foraging for themselves, also use vision to detect, approach and pursue live insects (**Figure 1)**. We previously showed that the house mouse, a popular model of mechanistic visual system development, readily approaches and ultimately preys on live insects using specific visual cues^13^, and that this behavior relies on specific neural circuitry in the superficial superior colliculus^25^. Here, we confirmed that P21 laboratory mice detect, approach and engage live crickets repeatedly in the laboratory setting (**Video 1**). We noted several important differences specifically in approach type orienting responses exhibited by P21 mice relative to mature adults when they first encounter live crickets. Most dramatically, P21 mice fail to show a binocular visual field bias upon initiating an approach relative to adults (**Figure 1A**). This correlates to a significant increase in the onset of the first approach towards a cricket by P21 mice versus adults (**Figure 1B**). Overall, these differences do not result in a decline in total number of approaches started over the entire 5 minute encounter with a live cricket (16.1 ± 1.2 vs. 17.5 ± 2.1, P21 vs. P90, N=12 vs. 10, respectively, p > 0.05, Welch’s t-test, **Video 1**). However, the noted detection and pursuit deficits do correlate with a significant decrease in the probability that an approach started will end with a successful contact (**Figure 1C & Video 1**). Further, when P21 mice successfully contacted live crickets, they did so from a wider mean stimulus angle at the end of approach (48.3 ± 2.2 vs. 26.1 ⨦ 0.5, P21 vs. P90, N=12 vs. 10, respectively, p < 0.01, Welch’s t-test) and engaged in prolonged proximate contact with rapid sniffing (**Figure 1D)**. Related to this behavior, juvenile mice rarely attacked insects even after repeated exposures over 5 days relative to adults (18.2% vs. 80% of juveniles vs. adults attacked crickets after 5 days of exposure for 5 minutes each day, Fisher’s Exact test, p = 0.0046, N=11 vs. 10, respectively), both groups without food deprivation.

**Figure 1.**
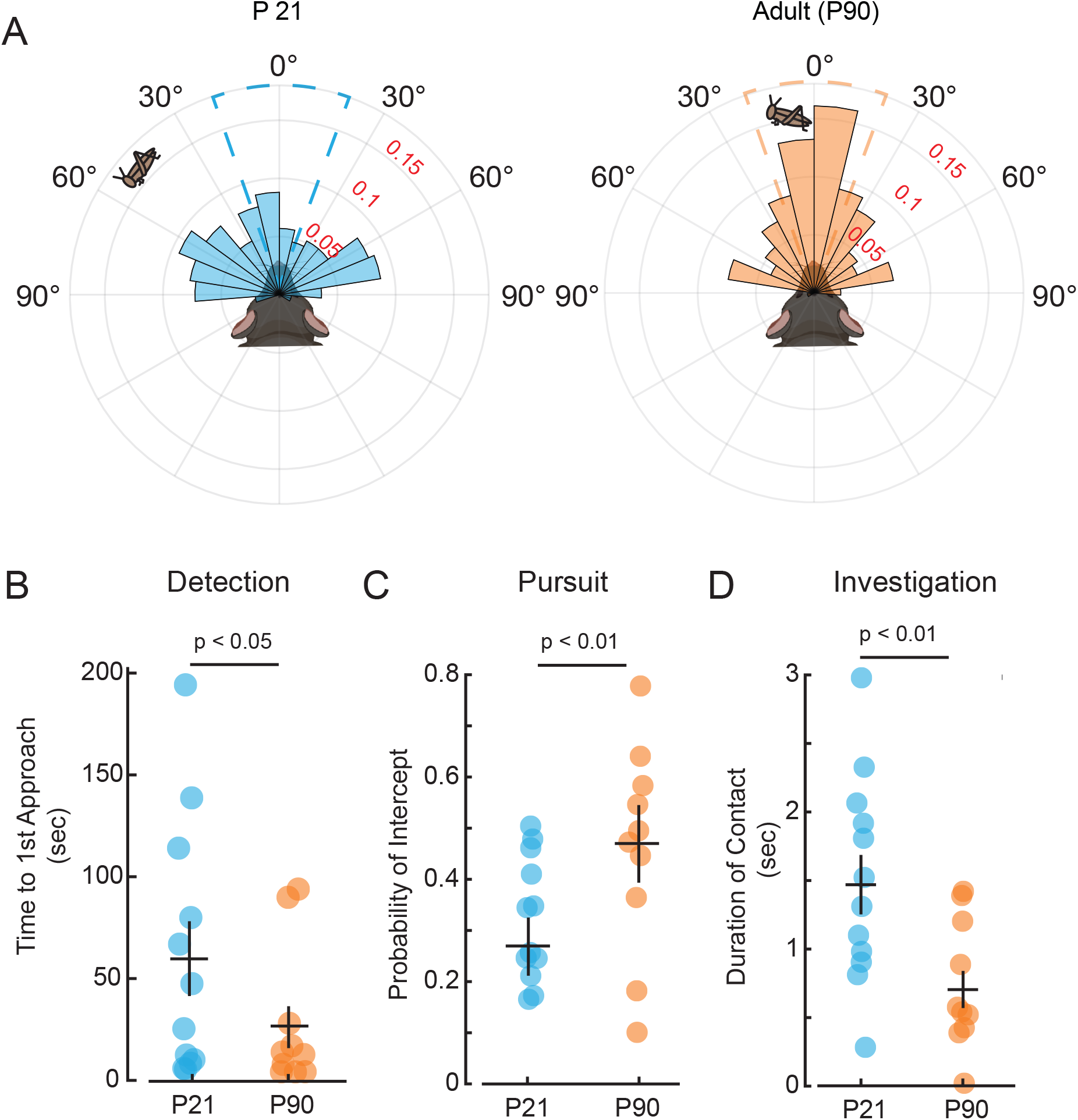
Analysis of natural prey capture behavior reveals a lack of binocular visual field bias to start approaches in P21 mice relative to adults. (A) Normalized polar plot distributions of stimulus angles (angle between mouse bearing and position of cricket) where successful approaches in juveniles (left) versus adults (right) began. (B) Detection behavior as measured by time to start an approach by P21 mice, blue, versus P90 mice, orange, (C) Probability of cricket interception, an approach that ends in cricket contact. (D) Stimulus investigation-related behavior indicated by the duration of a contact. Welch’s t-test, N=11 vs. 10, P21 mice vs. P90 mice, respectively, Error bars are +/-standard error of the mean.

Taken together, these results indicate that significant changes in visual field biases related to a mouse’s approach towards salient natural stimuli are immature in P21 mice. This developmental timepoint corresponds to known inabilities to integrate binocular information in that region of visual space as studied in visual cortex^1,28,29^ but could also relate to possible immaturity of complex binocular responses located within the adult mouse superior colliculus^9,30^. Consistent with this idea, induced perturbation of ipsilateral eye input to the superior colliculus in adults results in similar prey capture behavior deficits as we show characterize normally developing juveniles^24^. Our study therefore demonstrates for the first time, the perceptual consequences of previously identified immature binocular visual field processing in the mouse visual system.

Though live prey capture analysis captured a binocular processing deficit in developing mice, it is a complex multimodal experience relying on more sensory input than just vision^13^, especially in insect-naïve mice^31^. We therefore sought to uncover a more precise understanding of vision specific differences in particular that may be significantly different in the developing mouse. We therefore quantified the innate visual behavior of P21 mice evoked by virtual, high contrast motion stimuli displayed from a computer screen as we have done previously for adult mice^22^. We previously determined the features of simple moving ellipses that cause robust orienting responses in adult mice are more likely to elicit approach after they have experienced prey capture in the same environment^22^. This computerized-spontaneous perception of visual objects task, C-SPOT, affords the opportunity to vary specific parameters of visual objects and quantify how that alters stimulus detection, response, and egocentric sensory preferences (**Figure 2A-C**). We employed a simple version of this task by displaying a single objective stimulus size, shape, and speed of linear motion near the environmental horizon (**Figure 2A**) and found striking differences between P21 mice and mature adults in how they typically respond to the presented objective stimuli (**Figure 2D & E**). While both ages robustly approach the presented virtual stimulus over a total 5 minute presentation period, P21 mice took significantly longer to first approach (**Figure 2D & F**) and overall generated fewer approaches towards the stimulus over the entire session (**Figure 2D & G**). However, both P21 and adult mice are likely to arrest their locomotion in response to the presented stimuli with similar latencies from the start of presentation and display similar numbers of stimulus-evoked arrests (**Figure 2D, E, F & H**). This suggests that P21 mice are still capable of detecting the moving stimuli and find the motion salient but are specifically less likely to perceive the relative visual information that elicits an approach and allows it to end in a contact event. The increase in latency to approach the virtual targets by P21 mice is consistent in both our C-SPOT assay and live prey capture, suggesting that a difference in visual processing indeed explains the developmental differences in stimulus detection and specific aspects of pursuit quantified in the live prey capture assay.

**Figure 2.**
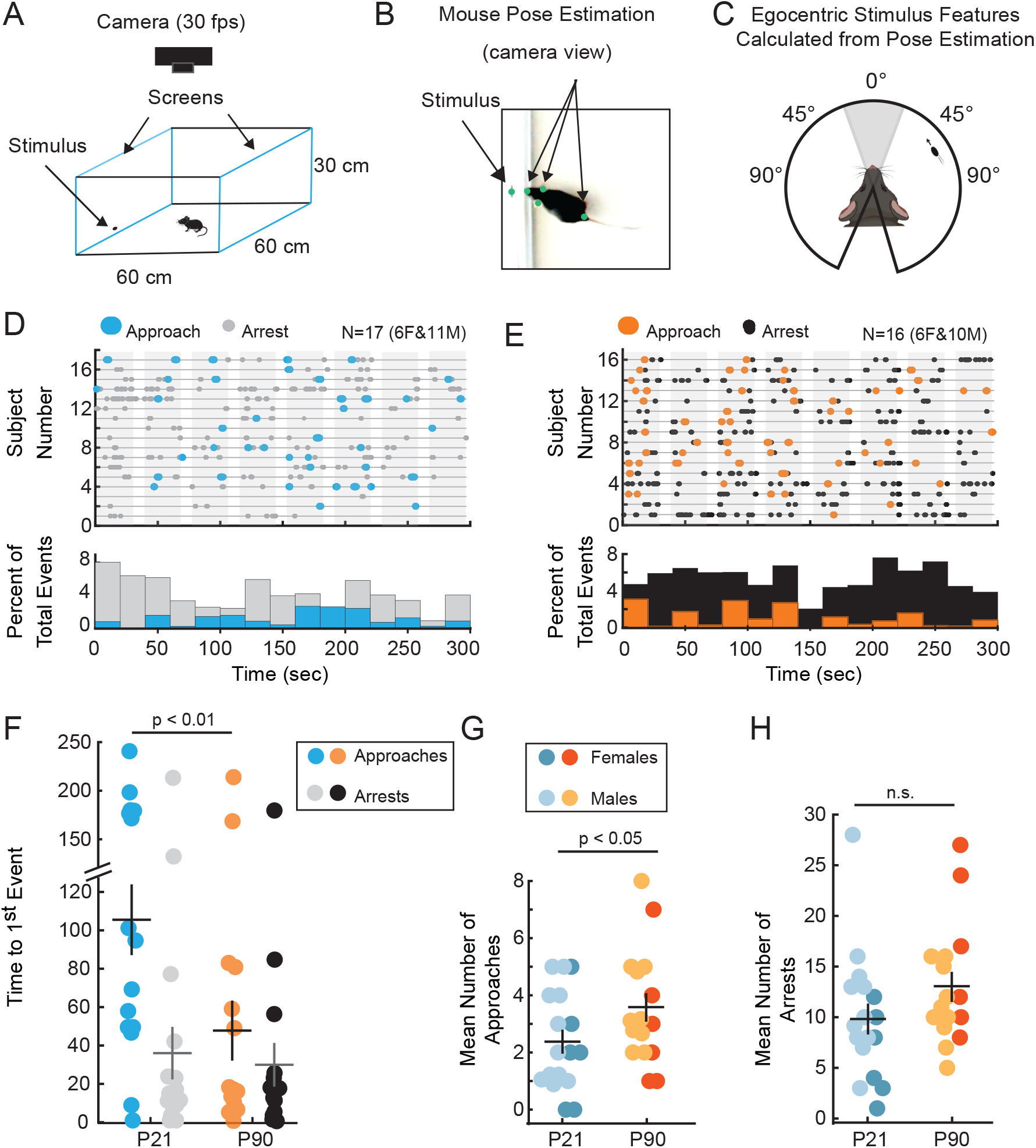
C-SPOT reveals robust developmental differences in innate visual orienting behavior. (A) Left, schematic of experimental arena. Blue outlines indicate location of computer monitors displaying stimuli and illuminating the environment. (B) Example frame from a recorded behavioral video overlaid with post estimates of relevant points. (C) Analysis of tracked positions in B, are used to generate estimates of egocentric visual features: relative stimulus size, speed, and position along the azimuth of the visual field (stimulus angle). (D & E) Ethograms of each subjects’ response (subject ID on y-axis) over time in seconds revealing when during each stimulus “sweep” (30 seconds, shaded in grey) approaches (color) or arrests (grey or black) occurred from stimulus onset. Ethograms are aligned to the start of 1^st^ stimulus presentation. Below, histograms showing the proportion of response type, either an approach (blue)or arrest (grey), that occurred during each 20 second bin of time. (F) Mean time to first orienting event of either approaches (colors) vs. arrests (grey and black). (G) Mean number of approach starts and (H) Mean number of arrests. Significance determined using Welch’s t-test, N = 17 vs. 16, P21 vs. P90 mice, respectively. n.s.=not significant. Error bars are +/-standard error of the mean.

To determine more precisely which visual responses and stimulus preferences are different between P21 mice and adult mice, we quantified the visual field location, relative size (in arc degree), and speed (in arc degree/sec) of the virtual stimulus at the onset of an approach or arrest response (**Figure 3**). Consistent with what was observed during live prey capture, P21 mice lack a strong binocular visual field bias in first detecting stimuli that they are able to successfully and continuously approach (**Figure 3A & B**, Left). The mean stimulus angle for when juvenile mice begin to approach a virtual moving target is further into the periphery, monocular visual field, around 60º (**Figure 3C**). In addition, even when P21 mice nearly contact the stimulus with the nose, they reach a position just in front of it whereas adults end with their nose touching at or near the center of the stimulus when they end an approach (**Videos 3 & 4)**. This leads to a significant difference in mean absolute stimulus angle at the end of an approach (33.1 ± 7.2 vs. 11 ± 4.3, P21 vs. P90, N=15 vs. 16, respectively, p < 0.05, Welch’s t-test). Taken together, these data demonstrate that P21 mice similarly detect and respond to stimuli located in their peripheral visual field, but specifically have a deficit in how they respond to stimuli that are located in central-anterior egocentric space, where visual information can be processed binocularly for adult animals.

**Figure 3.**
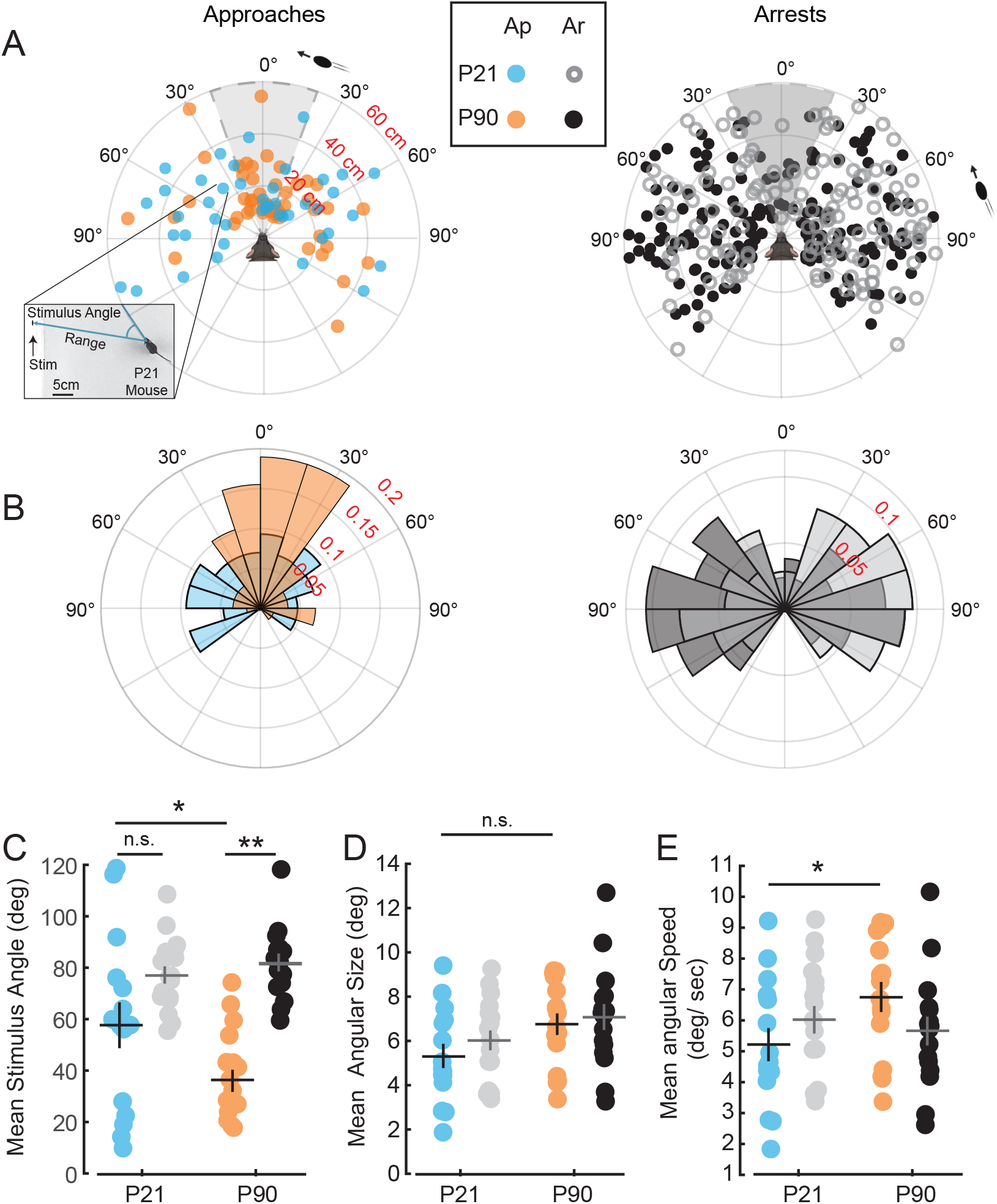
P21 mice show specific binocular visual field deficits at start of approach towards stimulus in the C-SPOT assay. (A) Polar plot distribution of individual approach starts (Left) and arrests (Right) evoked by a virtual visual stimulus for P21 (blue) versus P90 (orange) mice. Plotted is range (cm) versus stimulus angle (degree). Inset shows a frame from a P21 behavior video where an approach started towards a stimulus from a stimulus angle of ∼35 degrees from about 22cm away from the screen (represents blue point where call out to inset begins). (B) Normalized polar plots of approaches(Left) or arrests (Right) by age. Plotted is fraction of total events versus stimulus angle (degree). (C) Mean stimulus angle for each subject at approach starts (color) or arrests (black and grey). (D) Mean subjective size of stimulus (visual angle in degrees) when approaches (color) or arrests (black and grey) start. (E) Mean subjective speed of stimulus (degrees/sec) when approaches or arrests start. Significance determined using Welch’s t-test, with Benjamini-Hochberg procedure to correct for multiple comparisons, N = 17 vs. 16, P21 vs. P90 mice, respectively. n.s.=not significant, *= p< 0.05, **= p < 0.01. Error bars are +/-standard error of the mean.

Importantly, our assay revealed that the perception of stimuli that cause approach specifically are developmentally regulated relative to those perceptions that elicit arrest. For both ages, the stimuli that are likely to elicit arrest responses are primarily located in the periphery, but otherwise do not vary in relative size nor speed between juveniles and adults (**Figure 3D & E**). This suggests that the visual pathways that process peripheral, monocular visual information may have similar spatial and temporal resolution capabilities and lead to similar arrest behavioral outcomes at both ages. Indeed, we previously showed that the arrest responses evoked by these objective stimuli in the adults depend on the egocentric direction of motion of the stimulus across the visual field ^22^. That is, stimuli that elicited arrests not followed by an approach, were moving further into the peripheral monocular visual field of mice. In addition, both ages of mice were able to detect stimuli that caused either an approach (**Figure 3A**) or an arrest from the furthest possible reaches of the testing environment, ∼60cm from the mouse (**Figure 3**). Thus, P21 mice are capable of perceiving similar relative sizes and speeds of stimuli as compared to P90 adults. In contrast, P21 mice specifically show differences in their tendency to approach stimuli depending on the location of the stimulus within the visual field relative to P90 adults (**Figures 1,2 &3**). Importantly, we did not probe the capabilities of the juvenile visual system in this study exhaustively. We previously noted that receptive field properties do refine after eye-opening even in the monocular visual field of primary visual cortex ^32^. Thus, our future studies will seek to parametrically test a wider range of virtual stimulus sizes, speeds, and qualities of motion to further probe for differences in visual perception that change over development in retinotopic fashion.

To map developmental differences in mouse visual orienting behavior to their relevant larger scale neural networks, we assayed the expression of the immediate early gene c-Fos in the superior colliculus (**Figure 4**). We reasoned that this topographically organized, evolutionarily conserved, visual area may display significant difference in c-FOS activation as it has been significantly linked to generating representations of stimulus salience, direction of motion, relative size and speed of visual objects (see Basso et al.,^33^ for a recent review). Furthermore, specific cell types within the SC are required to control orienting behavior in the adult mouse during prey capture^25^ and the deeper layers that are aligned to the superficial retinotopy are known to integrate multimodal sensory input (visual, auditory and somatosensory) relevant to prey capture behavior^31,34,35^. We quantified the percent of c-Fos positive cells in different regions of the SC in mice exposed to C-SPOT versus age-matched controls exposed to the same environment with no visual stimulation (**Figure 4**). We obtained measures at three different locations along the anterior-posterior axis (-3.5, -4.15 and -4.6 AP), across three laminar zones from the dorsal surface (superficial, intermediate, and deep) and medial versus lateral divisions of the SC (**Figure 4A & B**). We calculated a ratio of c-Fos positive cells after visual stimulation normalized to age-matched controls without visual stimulation in each subregion of the SC. We then statistically compared those ratios between comparable subregions from the two ages of mice using Welch’s t-test with correction for multiple comparisons using the Benjamini-Hochberg procedure to decrease the false discovery rate as independence could not be assumed for within subject measures (i.e. AP, ML or depth position) (**Figure 4B**). This analysis revealed that c-Fos expression is significantly different between P21 and P90 mice within specific zones of the superior colliculus (**Figure 4B**, bolded entries). The anterior region of the superficial SC corresponds to the retinotopic location where stimuli fall on the ventral-temporal retina and thus occupy a relatively higher and central position in the animal’s visual field (**Figure 4A**). This region of the SC is involved in processing binocular input and distinct types of motion stimuli relevant to the orienting behaviors measured in our study^34–36^. Consistent with the relationship between the behavioral differences over development, c-Fos levels are significantly higher specifically in the anterior regions of the SC in the P21 mice exposed to virtual stimuli that cause approach and arrest behaviors. Although, it remains unclear why c-Fos levels are highly increased (two-to three-fold) in circuits that are maturing. One intriguing speculation for the regional specificity is that these early investigations of prey stimuli are engaging plasticity processes that upregulate cellular activity where changes in synaptic strength are needed. Future studies will seek to identify the specific cell types and circuits that upregulate c-Fos within the anterior SC in P21 mice and whether this expression reflects differences in neuronal activity measured during behavior or neuromodulation directed to this specific region of the superior colliculus.

**Figure 4.**
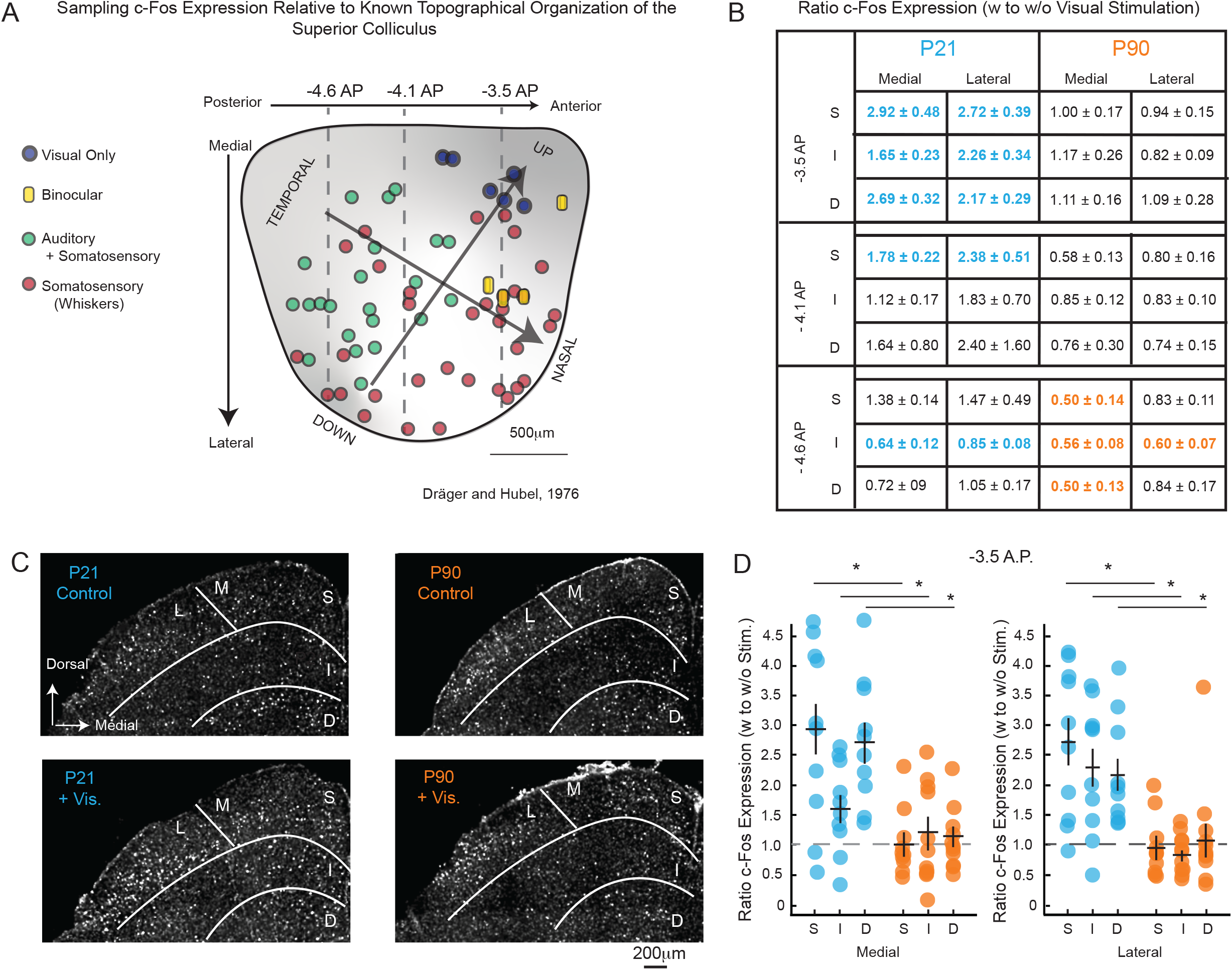
c-Fos expression differences after C-SPOT in P21 versus P90 mice. (A) Adaptation of response property mapping to the right hemisphere of the superior colliculus of the mouse from a dorsal view, Dräger and Hubel, 1976. The findings have been confirmed in subsequent decades (May, 2006) and led us to examine changes in c-Fos expression after exposure to virtual visual stimuli that evoke approach and arrest behaviors from three key planes of coronal section, -3.5 anterior-posterior (AP), -4.1 AP and -4.6 AP (grey dashed lines spanning the SC). Binocular visual processing is observed in the anterior SC (yellow rods) and overlays cells in the intermediate to deeper SC that respond to whisker stimulation near the nose (red circles underlying). We overlaid also the representation of discovered anisotropies in the representation of visual space across the surface of the SC, with darker shading of grey representing steeper changes in visual degrees over azimuth (temporal to nasal direction in the visual field) or elevation (Down to Up direction in the visual field). (B) Table summarizing the mean Ratio of c-Fos expression (percent of cells positive for c-Fos) in mice exposed to visual stimuli in C-SPOT relative to those w/out visual stimulation in the same environment. Ratios are normalized by dividing each subject’s percent of c-Fos positive cells by the mean of the percent of c-FOS positive cells in an age-matched control group, e.g., a ratio of 2.92 means a mouse after C-SPOT had nearly 3 times as many c-Fos positive cells in a given area than their age-matched control group only exposed to the environment for the same amount of time. Significant differences are highlighted in colored and bolded entries, P21 significant differences (p< 0.05) = blue and P90 (p< 0.05) significant differences =orange, N=10 and N=11. S= superficial, I= intermediate and D= deep, M= medial and L= lateral (C) Representative coronal sections of one hemisphere from each experimental group with quantified sub regions highlighted. +vis = samples from mice with visual stimulation during C-SPOT, Control = mice in arena for 10 minutes only, no displayed stimuli. (D) Example of normalized population data obtained from the medial and lateral regions of the superior colliculus from -3.5 AP from P21 mice (blue) versus P90 mice (orange). *=p<0.05, Welch’s t-test, followed by Benjamini-Hochberg procedure, N=10 and 11, P21 versus P90, respectively.

## Summary

Overall, we show significant developmental differences in how mice process stimuli in the binocular visual field that cause approach and pursuit behavior. It is noteworthy that these behavioral differences correlate with long-known significant developmental differences in visual receptive field properties in the primary binocular visual cortex^1-4^. However, our study finds important developmental differences in cellular activity during specific visual experiences within targeted regions of the SC that integrate binocular cues in the adult^9^. In previous studies of prey capture in mice where the pitch as well as the orientation of the head along the azimuth were quantified, it was found that mice angled their heads downwards during an approach which would position target stimuli higher up in the relative visual field^23^. A limitation of our current study is that we did not measure head pitch nor eye position in our animals and do not know whether there are developmental differences in these aspects of active visual sampling. However, previous work studying prey capture in adult mice which simultaneously measured eye positions during prey capture has shown that the head angle of the mouse in the azimuth can be used to approximate the position of the binocular visual field^23,37^. Therefore, the evidence in our study argues for more targeted studies of visual stimulus encoding development and plasticity in the superior colliculus, with an emphasis on understanding the emergence and plasticity of the diverse forms of binocular processing recently quantified in the SC^9^.

Our study also demonstrated that C-SPOT may be used on young, developing mice to quantify complex visual behavior driven by visual stimulation only. This will be of significant use for those requiring a behavioral tool to assess models of neurodevelopmental disorders, Alzheimer’s, schizophrenia or other diseases that impact sensory stimulus detection, selection and visual orienting. Indeed, Autism-associated Shank3 knockout in mice leads to robust deficits in ocular dominance plasticity^38^. It will be interesting to determine whether our assay can more precisely quantify the nature of visual perception deficits in these mice and how interventions to restore gene and neural circuit function in these mice may impact their visual perception through development.

Finally, C-SPOT itself may be easily modified to test more fundamental qualities of natural visual processing in the mouse as has been highly developed for the larval zebrafish^38,39^. This would provide a bridge between past and future groundbreaking discoveries made in the larval zebrafish to the principles underlying mammalian visual system development and function^40^. In addition, the clear behavioral readouts that we quantify with C-SPOT will lead to the development of many creative visual “illusions” to test the limits and capabilities of mouse visual perception using a “naturalistic behavior screening” approach to complement and expand on hypotheses-based testing of our limited knowledge of the receptive field properties so far observed in the visual system of the mouse^34,41–45^.

## Materials and Methods

### Subjects

All animals were used in accordance with protocols approved by the University of Nevada, Reno, Institutional Animal Care and Use Committee, in compliance with the National Institutes of Health Guide for the Care and Use of Laboratory Animals. Both male and female mice were used in the study. We tested 29 P20-22 juvenile mice and 26 adult mice, aged P90 or greater; the specific number used in each statistical comparison is noted in figure legends and results. We used C57BL/6J and mixed background transgenic mice: Ntsr1-GN209-Cre and Grp-KH288-Cre lines^46^. Mice were group housed, up to 5 animals per cage excluding single housed mice from the study. Mice had ad libitum access to water and food (Envigo, Teklad diet, 2919). The vivarium was maintained on a 12 hour light/dark schedule, and all behavioral measures were acquired within 3 hours of the dark to light transition.

### Apparatus

Mice were individually removed from their home cages and placed into a rectangular, white acrylic open field arena with aluminum slot framing (McMaster-Carr), 60 cm long x 60 cm wide x 30 cm high, with 1/32” thick white buna-N /vinyl rubber flooring (McMaster-Carr) (**Figure 2A**). A pair of opposing sides of the arena consisted of Hewlett Packard VH240a video monitors that measured 60.5 cm diagonally, with a vertical refresh rate of 50-60 Hz and resolution set to 1920×1080 pixels. A solid white background was displayed on both monitors to achieve even lighting throughout the arena. Visual stimuli could then appear on either monitor in C-SPOT trials. Crickets during live prey capture trials were placed into the arena by the experimenter at a location away from the mice. A Logitech HD Pro Webcam C920 digital camera was suspended overhead to capture the behavior at 30 frames per second throughout each trial.

### Visual Stimuli

Visual stimuli for C-SPOT were generated with MATLAB Psychophysics toolbox (Brainard, 1997) and displayed on an LED monitor (60 Hz refresh rate, ∼50 cd/m2 luminance) in a dark room. To mimic insect proportions, we displayed ellipses with a major axis that was 2 times the size of the minor axis and displayed stimuli that were 2 cm along the horizontal axis as these stimuli evoked the most frequent approaches in adult mice^22^. For a presented stimulus with a major axis of 2 cm, this corresponds to the presentation of a ∼4 degree sized target from 30 cm away from the screen. Stimulus speed was kept constant at 2 cm/sec as this speed evoked the most approach and least amount of arrests in adult mice as previously reported in Procacci et al., 2020. This meant that a single stimulus sweep lasted for approximately 30 sec on the screen, with the stimulus disappearing for approximately 7 seconds before reappearing for the next 30 sec sweep. This pattern lasted the extent of the trial, 300 seconds.

### Behavior

As described previously, Procacci et al., 2020, prior to C-SPOT or live prey capture, mice were acclimated to handlers and arena for 4 days during which each mouse was handled 3 times a day for 3 minutes each and placed in the arena 3 times a day for 5 minutes each time. Young mice were habituated to handler and environment 3-4 days prior to their tested age of P20-22. On the testing/behavioral measurement day, group housed mice were brought into the testing room in their home cages and allowed to acclimate to the darkened testing room. Then, each experimental subject was placed in the testing arena and habituated for 3-5 minutes, with controlled illumination emitted from computer screen monitors only. Either C-SPOT was run, or live crickets were introduced and behavior recorded. The arena floor was cleaned thoroughly with 70% EtOH after each mouse was removed to mitigate odor distractions. Exposing mice to stimuli did not begin until each mouse demonstrated self-grooming behaviors and reduced defecation and thigmotaxic behavior which were taken as indication they were not anxious in the environment, again, around 3-5 minutes.

For testing prey capture behavior, mice were given 10 minutes with the cricket on each day after habituation for up to 5 days and were not food deprived in order to understand how visual stimuli were interpreted at an individual’s baseline state. If mice caught the cricket within this period of time, they were given another until up to three were caught, after which mice were returned to their home cage. If mice did not catch the cricket within 10 minutes, the experimental subject was returned to their home cage, the arena floor was cleaned, and another mouse was given a chance. All mice were then returned to their home cages with standard food. This protocol was repeated for 5 days, until most adult mice reliably caught crickets in the arena in the 10 minute period. The crickets used were *Acheta domestica* obtained from Fluker’s Farm or a local pet store and were 1-2 cm in length, group-housed, and fed Fluker’s Orange Cube Cricket Diet.

### Data Analysis

DeepLabCut was used to digitize and extract 2-dimensional coordinates of the mouse’s nose, two ears and body center, as well as the center point of the stimulus (cricket or computer-generated stimulus) throughout the video recordings at 30Hz. These tracks were entered into customized MATLAB scripts to extract behavioral parameters: mouse speed, stimulus speed, stimulus angle, range, subjective stimulus size and speed.

An arrest “response” was defined as any time the mouse’s nose and body moved less than 0.5 cm/sec for a duration of 0.5 -2 seconds. Arrests that occurred in the absence of a visual stimulus, or when the stimulus was more than 140 degrees from the bearing of the nose, were excluded from analysis of visual driven arrest responses. Approach starts were defined as mice moving toward the stimulus starting from a distance of at least 8 cm, and at an average approach speed of at least 15 cm/sec and a stimulus bearing of less than 150 degrees. Using these definitions, we computed the percentage of stimulus trials in which each behavior was observed, as well as the number of arrests and approaches that occurred during individual trials. A successful approach was defined as any time the mouse’s nose came within 3 cm of the stimulus after an approach start had been identified.

For both response types, approach or arrest, we calculated range as the distance between the center of the head between the two ears and the center of the stimulus, and stimulus angle as the angular distance between the line emanating from the center of the mouse body to the mouse nose, and that from the center of the mouse body to the center of the stimulus. Angular stimulus sizes and speeds were calculated using the horizontal length of the stimulus (2 cm) and the distance of the behavioral event from the stimulus in cm.

Statistics were performed using MATLAB and R software. Where means are reported and data are normally distributed we used Welch’s t-test (two group comparisons), followed by Benjamini-Hochberg procedure to decrease false discovery rate and correct for multiple comparisons. The specific tests used are specified in figure legends. Where medians are reported and/or data are not normal, rank sum tests were used. Significant differences in proportional measures were determined by Fisher’s exact test. Test results with a p-value of < 0.05 were considered significant. Cohen’s D > 0.8 considered a large effect, with all significant effects in this study having Cohen’s D > 0.8.

### Immunohistochemistry

To assay c-Fos expression in mice exposed to C-SPOT and age-matched controls without visual stimulation and experience, mice were deeply anesthetized ninety minutes after behavioral testing through inhalation with 3.5% isoflurane. Subjects were then transcardially perfused with 75 ml of phosphate buffered saline (1X PBS) followed with 25 ml of 4% paraformaldehyde (PFA). Brains were removed and stored in 4% PFA for 24 hours at 4°C. Brains were then removed from PFA and rinsed with PBS before sectioning into 45 μm thick coronal sections. Floating sections were stored in 0.02% sodium azide (NaN_3_) and PBS at 4°C for up to two weeks before assaying for c-Fos protein expression. To assay c-Fos protein expression, floating sections in well dishes were incubated in a blocking solution (4% bovine serum albumin (BSA), 2% horse serum, 0.2% Triton X-100, 0.05% NaN_3_) for 3 hours, followed by incubation with rabbit anti-c-Fos primary antibody (1:500, 226 003, Synaptic Systems) in blocking solution for 24 h at 4°C. Sections were washed 3 times for 20 minutes each time in PBS at 20°C before incubating with goat anti-rabbit Alexa 488 secondary antibody (1:500, A32723 Invitrogen) in blocking solution for 2 h at 20°C. Sections were then incubated in DAPI (1:1000, 64427, ThermoFischer Scientific; 15 min at 20°C) diluted in PBS for nuclei labeling. Sections were washed again 3 times for 20 minutes each time before mounting onto slides with AquaPoly mounting media (18606, Polysciences). Mounted sections were stored at 4°C in dark slide boxes until image processing.

### c-Fos Quantification

Images were obtained using a Leica Confocal microscope. Tile scans of the entirety of each coronal slice was imaged with a 20X objective and saved as a LIF image file. Sections between -3.5 to -4.59 AP were imaged. Specific regions and structures were identified in as in Paxinos and Franklin’s the Mouse Brain in Stereotaxic Coordinates as well as the Allen Mouse Brain Coronal Atlas (https://mouse.brain-map.org/static/atlas). Subregions and laminae of colliculus were identified as in Wang and Burkhalter, 2013^47^.

c-Fos positive cells were identified and quantified using Imaris Microscopy Image Analysis Software (Oxford Instruments). LIF files were loaded into IMARIS and Gamma correction (.8%) and background subtraction were standardized and applied similarly to all images. Images were analyzed blind to testing condition. Three circular regions of interest (ROIs) were drawn in each collicular layer: superficial, intermediate, and deep in both the medial and lateral regions. To ensure similarity of sub regions quantified between subjects, ROIs for the medial region spanned the center of a region as measured from the brain’s midline (medial border) to the medial/lateral border of the section (halfway between the medial-lateral borders of the SC, **Figure 4C**). Lateral ROIs were drawn halfway from the medial/lateral border of the section to the lateral border of the SC. The number of c-Fos positive cells and DAPI positive cells for each ROI were used to create a ratio of c-Fos:DAPI fluorescence. These percentages were then averaged per medial or lateral region, and again averaged between 2-3 sections for each AP region for each subject. These “per subject” percent averages were then used to determine the relative increase in c-Fos positive cells between visually-stimulated mice and their controls (age-matched and exposed to environment only). The reported normalized metric “ratio of c-Fos expression (w to w/o)” is each experimental subject’s percent of c-Fos positive cells divided by the average percentage obtained from their age-matched controls that were placed in the same environment with no visual stimulation. This therefore compares the percent increase in c-Fos positive cells when mice view and respond to visual stimuli in our testing arena, versus when mice wander in the same arena without the specific visual experience assayed.

### Author Contributions

K.A. collected and analyzed the behavioral and histological data and co-wrote and edited the manuscript. R.G.O. collected and analyzed the live prey capture behavior data. M.K. analyzed histological data. T.F. and S.P. developed the histological approaches used in the manuscript and provided input on experimental design and drafts of the manuscript. J.L.H devised the experiments, coordinated and obtained funding for the study, analyzed the behavioral and histological data, co-wrote and edited the manuscript.

## Supporting information

Video 1 P21 Cricket

Video 2 Adult Cricket

Video 3 P21 Virtual Stimulus

Video 4 Adult Virtual Stimulus

## Acknowledgements

We thank Anastasiya Chevychalova for manually scoring behaviour. We greatly appreciate input on drafts of our manuscript from Drs. Jianhua Cang, David Feldheim, Jason Triplett, Angie Michaiel, Philip Parker and Adema Ribic.

## References

1. Hensch, T.K. Annu. Rev. Neurosci. 27, 549–579 (2004).

2. Cang, J., Savier, E., Barchini, J. & Liu, X. Annu Rev Vis Sci 4, 239–262 (2018).

3. Leighton, A.H. & Lohmann, C. Front. Neural Circuits 10, 71 (2016).

4. Hooks, B.M. & Chen, C. Neuron 56, 312–326 (2007).

5. Tan, L., Tring, E., Ringach, D.L., Zipursky, S.L. & Trachtenberg, J.T. Neuron 108, 735– 747.e6 (2020).

6. Tan, L., Ringach, D.L. & Trachtenberg, J.T. J. Neurosci. (2022).doi:10.1523/JNEUROSCI.1702-21.2022

7. May, P.J. Progress in Brain Research 321–378 (2006).doi:10.1016/s0079-6123(05)51011-2

8. Ito, S. & Feldheim, D.A. Front. Neural Circuits 12, 10 (2018).

9. Russell, A.L., Dixon, K.G. & Triplett, J.W. Journal of Neurophysiology 127, 913–927 (2022).

10. Wheatcroft, T., Saleem, A.B. & Solomon, S.G.doi:10.31234/osf.io/2yaj7

11. Storchi, R. et al. Sci. Rep. 9, 10396 (2019).

12. El-Danaf, R.N. & Huberman, A.D. Journal of Comparative Neurology 527, 259–269 (2019).

13. Hoy, J.L., Yavorska, I., Wehr, M. & Niell, C.M. Curr. Biol. 26, 3046–3052 (2016).

14. Dean, P., Redgrave, P. & Westby, G.W.M. Trends in Neurosciences 12, 137–147 (1989).

15. Ngo, V. et al.doi:10.1101/2022.04.01.486794

16. Klomp, H. & Tinbergen, L. Archives Néerlandaises de Zoologie 13, 344–379 (1960).

17. Borghuis, B.G. & Leonardo, A. J. Neurosci. 35, 15430–15441 (2015).

18. Lin, H.-T. & Leonardo, A. Curr. Biol. 27, 1124–1137 (2017).

19. Roulin, A. (Cambridge University Press: 2020).

20. Spinka, M., Newberry, R.C. & Bekoff, M. The Quarterly Review of Biology 76, 141–168 (2001).

21. Oldfield, C.S. et al. Elife 9, (2020).

22. Procacci, N.M. et al. Proc. Biol. Sci. 287, 20201189 (2020).

23. Johnson, K.P. et al. Neuron 109, 1527–1539.e4 (2021).

24. Su, J. et al. Proceedings of the National Academy of Sciences 118, (2021).

25. Hoy, J.L., Bishop, H.I. & Niell, C.M. Curr. Biol. 29, 4130–4138.e5 (2019).

26. Igelstrom, K.M., Herbison, A.E. & Hyland, B.I. Neuroscience 168, 706–714 (2010).

27. Tatavarty, V. et al. Neuron 106, 769–777.e4 (2020).

28. Chang, J.T., Whitney, D. & Fitzpatrick, D. Neuron 107, 338–350.e5 (2020).

29. Wang, B.-S., Sarnaik, R. & Cang, J. Neuron 65, 246–256 (2010).

30. Hoffmann, K.P. & Sherman, S.M. Journal of Neurophysiology 38, 1049–1059 (1975).

31. Shang, C. et al. Nat. Neurosci. 22, 909–920 (2019).

32. Hoy, J.L. & Niell, C.M. J. Neurosci. 35, 3370–3383 (2015).

33. Basso, M.A., Bickford, M.E. & Cang, J. Neuron 109, 918–937 (2021).

34. Drager, U.C. & Hubel, D.H. Nature 253, 203–204 (1975).

35. Drager, U.C. & Hubel, D.H. Journal of Neurophysiology 39, 91–101 (1976).

36. de Malmazet, D., Kühn, N.K. & Farrow, K. Curr. Biol. 28, 2961–2969.e4 (2018).

37. Michaiel, A.M., Abe, E.T. & Niell, C.M. Elife 9, (2020).

38. Tatavarty, V. et al.doi:10.1101/365445

39. Barker, A.J. & Baier, H. Curr. Biol. 25, 2804–2814 (2015).

40. Zimmermann, M.J.Y. et al. Curr. Biol. 28, 2018–2032.e5 (2018).

41. Marshel, J.H., Garrett, M.E., Nauhaus, I. & Callaway, E.M. Neuron 72, 1040–1054 (2011).

42. Juavinett, A.L. & Callaway, E.M. Curr. Biol. 25, 1759–1764 (2015).

43. Esfahany, K., Siergiej, I., Zhao, Y. & Park, I.M. eNeuro 5, (2018).

44. Barchini, J., Shi, X., Chen, H. & Cang, J. eLife 7, (2018).

45. Lee, K.H., Tran, A., Turan, Z. & Meister, M. eLife 9, (2020).

46. Gerfen, C.R., Paletzki, R. & Heintz, N. Neuron 80, 1368–1383 (2013).

47. Wang, Q. & Burkhalter, A. Journal of Neuroscience 33, 1696–1705 (2013).

